# Unlocking the Potential: FKK6 as a Microbial Mimicry-Based Therapy for Chronic Inflammation-Associated Colorectal Cancer in a Murine Model

**DOI:** 10.1101/2024.07.30.605845

**Authors:** Lucia Sládeková, Hao Li, Vera M. DesMarais, Amanda P. Beck, Hillary Guzik, Barbora Vyhlídalová, Haiwei Gu, Sridhar Mani, Zdeněk Dvořák

## Abstract

Chronic intestinal inflammation significantly contributes to the development of colorectal cancer (CRC) and remains a pertinent clinical challenge, necessitating novel therapeutic approaches. Indole-based microbial metabolite mimics FKK6, which is a ligand and agonist of the pregnane X receptor (PXR), was recently demonstrated to have PXR-dependent anti-inflammatory and protective effects in a mouse model of dextran sodium sulfate (DSS)-induced acute colitis. Here, we examined the therapeutic potential of FKK6 in a mouse model (C57BL/6 FVB humanized PXR mice) of colitis-associated colon cancer (CAC) induced by azoxymethane (AOM) and dextran sodium sulfate (DSS). FKK6 (2 mg/kg) displayed substantial anti-tumor activity, as revealed by reduced size and number of colon tumors, improved colon histopathology, and decreased expression of tumor markers (c-MYC, β-catenin, Ki-67, cyclin D) in the colon. In addition, we carried out the chronic toxicity (30 days) assessment of FKK6 (1 mg/kg and 2 mg/kg) in C57BL/6 mice. Histological examination of tissues, biochemical blood analyses, and immunohistochemical staining for Ki-67 and γ-H2AX showed no difference between FKK6-treated and control mice. Comparative metabolomic analyses in mice exposed for 5 days to DSS and administered with FKK6 (0.4 mg/kg) revealed no significant effects on several classes of metabolites in the mouse fecal metabolome. Ames and micronucleus tests showed no genotoxic and mutagenic potential of FKK6 *in vitro*. In conclusion, anticancer effects of FKK6 in AOM/DSS-induced CAC, together with FKK6 safety data from *in vitro* tests and *in vivo* chronic toxicity study, and comparative metabolomic study, are supportive of the potential therapeutic use of FKK6 in the treatment of CAC.

## INTRODUCTION

Inflammatory bowel disease (IBD), encompassing Crohn’s disease and ulcerative colitis (UC), is a chronic condition characterized by a well-established correlation with colorectal cancer (CRC). Patients diagnosed with IBD face 2- to 8-fold elevated risk of developing CRC [1]. This remarkable association underscores the deep connection between chronic inflammation, such as colitis, and the initiation of colitis-associated colon cancer (CAC). Chronic inflammation plays a crucial role in carcinogenesis by creating a pro-tumorigenic microenvironment that promotes cell proliferation, leads to mutations, and induces genomic instability [2, 3] and epigenetic changes that silence tumor suppressor genes and activate oncogenes [4]. Therefore, attenuating intestinal inflammation is one of the strategies for preventing CAC onset in IBD patients. Various inflammation-related cellular objects were proposed or employed as the pharmacological targets for a chemoprevention of CAC. The examples, but not exhaustive list, comprise antibodies against inflammatory cytokines (Adalimumab, Infliximab, Tocilizumab) and cytokines receptors (Basiliximab, Daclizumab), selective cyclooxygenase 2 (COX-2) inhibitors (Celecoxib, Rofecoxib), COX-1/2 inhibitors (aspirin), and various non-steroidal anti-inflammatory drugs (NSAID) [5]. For instance, long-term users of aspirin and other NSAIDs had reduced incidence of CRC by 40-50% [6]. 5-aminosalicylic acid was associated with a 49% reduced risk of adenoma colorectal neoplasms in UC patients. European Cancer Organization recommended the use of mesalamine compounds (aminosalicylates) to UC patients [1].

An emerging and promising therapeutic target in the treatment of CAC could be the pregnane X receptor (PXR). Besides its central role in the regulation of xenobiotic-metabolizing enzymes, the PXR controls the intermediary metabolism of lipids, glucose, and bile acids, and it is involved in the onset and progress of various diseases, including diabetes, metabolic syndrome, cardiovascular pathologies, acute kidney injury, neurological pathologies, inflammatory bowel disease, and cancer. Multiple *in vivo* studies revealed the functions of PXR in IBD. Attenuated activation of the PXR by endogenous bile acids [7] and decreased expression of the PXR in the intestines were reported in patients with Crohńs disease [8]. Consistently, the amelioration of chemically induced colitis *in vivo* was achieved by various PXR agonists such as flavonoids [9] and sesquiterpenes [10] by the NFκB inhibition mechanism.

On the other hand, the roles of PXR in CRC are context-dependent. PXR accounts for reduced effectivity of some CRC chemotherapeutics, such as doxorubicin [11] and irinotecan [12]. PXR upregulated anti-apoptotic genes [13], de-repressed p53 [14], and enhanced the neoplastic characteristics [15] in human colon cancer cells. In addition, PXR was identified as a key factor for colon cancer stem cells self-renewal and chemoresistance [16]. Therefore, PXR is a potential biomarker of unfavorable outcomes in CRC chemotherapy or radiation [12, 13, 15–28]. On the other hand, in a single study, PXR suppressed the proliferation and tumorigenicity of colon cancer cells by controlling the cell cycle in endogenously low PXR-expressing cells [29]. The activation of PXR by rifaximin protected against azoxymethane (AOM)/dextran sulfate sodium (DSS)-induced colon cancer by the mechanism involving blockade of NFκB [30]. Rifaximin also inhibited the release of pro-angiogenic mediators in colon cancer cells through the PXR-dependent pathway [31]. Collectively, the PXR roles in intestinal pathophysiology resemble Dr. Jekyll and Mr. Hyde’s story. The appropriate activation of PXR by endogenous, microbial, and dietary ligands and the physiological expression of PXR in the gut are essential conditions for intestinal health.

On the contrary, decreased expression of PXR, its insufficient endogenous activity, or its excessive activation by xenobiotics are associated with the onset and progress of IBD and CRC (CAC). [32, 33]. Recently, we have demonstrated a novel concept of small-molecule mimicry of microbial metabolites present in humans to expand drug repertoires [34]. We designed a series of highly potent, efficacious, non-toxic PXR-selective agonists mimicking indole-based intestinal microbial metabolites. Lead derivative FKK-6 displayed PXR-mediated *in vitro anti-inflammatory and in vivo* anti-colitis phenotype [35]; however, the effects in colitis-induced colon cancer is unknown.

Recently, we reported on the initial *in vitro* pharmacological profiling of FKK6. High selectivity of FKK6 towards PXR was supported by hydrogen-deuterium exchange mass spectrometry and broad counter-screen against potential off-targets, including G protein-coupled receptors, steroid and nuclear receptors, ion channels, and xenobiotic membrane transporters. Solubility in simulated gastric and intestinal fluids, the partition coefficient, and permeability in Caco2 cells were indicative of essentially complete *in vivo* absorption of FKK6. It was rapidly metabolized in human liver microsomes, yielding two metabolites with significantly reduced PXR agonist potency, which implies that despite high intestinal absorption, FKK6 is eliminated quickly by the liver, and its PXR effects are predicted to be predominantly in the intestines. FKK6 has exhibited a suitable pharmacological profile supporting its potential preclinical development [36].

In the current study, FKK6 is evaluated for its potential to abrogate tumor number and size in the AOM-DSS mouse model of colitis-induced colorectal cancer. In addition, chronic toxicity studies were conducted in mice, and fecal metabolomics was assessed in mice exposed to DSS for 5 days.

## MATERIALS AND METHODS

### Chemicals and reagents

Acetonitrile (ACN; A998-1), methanol (MeOH; A452-4), 10× PBS solution (50-105-5502), and acetic acid (A38C-212), all LC-MS grade, were purchased from Fisher Scientific (Pittsburgh, PA). Ammonium hydroxide (AX1308-7), ammonium acetate (A-1542) was bought from Sigma-Aldrich (Saint Louis, MO). De-ionized water was provided in-house by a Water Purification System from EMD Millipore (Billerica, MA). PBS was bought from GE Healthcare Life Sciences (Logan, UT). FKK6 was synthesized as described previously [35]. The standard compounds corresponding to the measured metabolites were purchased from Sigma-Aldrich (Saint Louis, MO) and Fisher Scientific (Pittsburgh, PA).

### Ames fluctuation test and bacterial cytotoxicity

Ames fluctuation test was performed in *Salmonella typhimurium* strains TA98, TA100, TA1535, and TA1537 in the presence or absence of rat liver S9 fraction [37]. Bacteria were incubated for 96 h at 37 °C with FKK6 (5 μM – 100 μM), quercetin (QUE; 30 μM), streptozotocin (STR; 2.5 μM), 9-aminoacridine (9AA; 10 μM), 2-aminoanthracene (2AA; 10 μM), Mitomycin C and vehicle. The cytotoxicity results were expressed as a percent of control growth (OD_650_). Compounds with the effects on the growth lower than 60% of control were considered cytotoxic. Wells that displayed bacteria growth due to the reversion of the histidine mutation (as judged by the ratio of OD_430_/OD_570_ being greater than 1.0) were counted and recorded as positive counts. The significance of the positive counts between the treatments and the controls was calculated using the one-tailed Fisher’s exact test. The significance levels were applied as follows: Weak positive (“+”), if 0.01 ≤ *p* < 0.05; Strong positive (“++”), if 0.001 ≤ *p* < 0.01; Very strong positive (“+++”), if *p* < 0.001.

### *In vitro* micronucleus assay

Dihydrofolate reductase deficient Chinese hamster ovary cells (CHO-K1) were incubated at 37°C with FKK6 (8 μM; 16 μM; 32 μM), Mitomycin C (MTC; 0.3 μM), cyclophosphamide (CP; 72 μM) or vehicle (DMSO; 1%) in the presence (4 h) or absence (24 h) of rat liver S9 fraction, as described elsewhere [38]. The percent of micronucleated cells was calculated. A marginally positive result (“-/+”) was defined as a value significantly higher than controls (t-test, p < 0.05), and at least 2-fold higher than controls. A positive result (“+”) is defined as a value significantly higher than controls (t-test, p < 0.05) and at least 3-fold higher than controls.

### Metabolomics

The targeted LC-MS/MS method was used in numerous studies [39–41]. LC-MS/MS experiments were performed on an Agilent 1290 UPLC-6490 QQQ-MS (Santa Clara, CA) system. Each sample was injected twice: 10 µL for analysis using negative ionization mode, and 4 µL for analysis using positive ionization mode. Chromatographic separations were carried out in hydrophilic interaction chromatography (HILIC) mode on a Waters XBridge BEH Amide column (150 × 2.1 mm, 2.5 µm particle size, Waters Corporation, Milford, MA). The flow rate was 0.3 mL/min, the auto-sampler temperature was kept at 4 °C, and the column compartment was set at 40 °C. The mobile phase was composed of Solvents A (10 mM ammonium acetate, 10 mM ammonium hydroxide in 95% H_2_O/5% ACN) and B (10 mM ammonium acetate, 10 mM ammonium hydroxide in 95% ACN/5% H_2_O). After the initial 1 min isocratic elution of 90% B, the percentage of Solvent B decreased to 40% at t=11 min. The composition of Solvent B was maintained at 40% for 4 min (t=15 min), and then the percentage of B gradually went back to 90% to prepare for the next injection. The mass spectrometer is equipped with an electrospray ionization (ESI) source. Targeted data acquisition was performed in multiple-reaction monitoring (MRM) mode. The whole LC-MS system was controlled by Agilent Masshunter Workstation software (Santa Clara, CA). The extracted MRM peaks were integrated using Agilent MassHunter Quantitative Data Analysis (Santa Clara, CA).

### Animals

Five-week-old male or female C57BL/6 mice (Jackson Laboratories, Bar Harbor, Maine; # 000664) were co-housed for acclimatization at the vivarium for 2 weeks before experiments. For certain experiments like the AOM-DSS study, which faithfully replicates human colorectal cancer genomics [42], FVB humanized PXR mice (∼67% FVB/N)(see genotyping details from DartMouse in Zenodo: http://doi.org/10.5281/zenodo.12187327) were used (kindly provided by Julia Cui, Washington University, St. Louis). All animal experiments were approved by the Animal Institute Committee (Protocols #00001409 & #00001610) of the Albert Einstein College of Medicine and were performed following institutional and national guidelines. All mice were sex and age-matched within experiments and maintained under a strict 12-hour light/dark cycle with free access to sterilized chow and water.

#### Chronic toxicity assessment (30 days)

Seven-week-old C57BL/6 mice were divided into three groups – 6 mice in each group with 3 females and 3 males. Group #1: control (100 μL of normal saline with 1% DMSO); Group #2: FKK6 – 1 mg/kg (100 μL of 0.5 mM FKK6); Group #3: FKK6 – 2 mg/kg (100 μL of 1 mM FKK6). Stock solutions of test compound were diluted in normal saline. Mice were orally gavaged daily for a period of 30 days. On Day 30, mice were euthanized, and sera were collected for blood biochemistry assays - Superchem Analysis (SA020, Antech Diagnostics, New Hyde Park, NY). Tissues were collected by Albert Einstein Histology and Comparative Pathology Facility and the histology was evaluated at Albert Einstein Histology and Comparative Pathology Facility. Ki-67 immuno-histochemical staining of gastro-intestinal sections, including both small and large intestines, and data analyses were done at Albert Einstein College Analytical Image Facility. The same tissue sections were stained and analyzed for the expression of γ-H2AX (phospho Ser139) (HistoWiz, Inc., New York City, NY, USA).

#### AOM Assay

6-8 weeks FVB humanized PXR mice were administrated with a single i.p. dose of 10 mg/kg AOM (Azoxymethane, A5486, Sigma) on Day 1, then treated with 3% (w/v) DSS in water (molecular weight, 36,000–50,000; MP Biomedicals Inc.) for 7 days. After 7 days, 3% DSS was replaced by regular water for 7 days. The cycle (7 days 3% DSS and 7 days regular water) was repeated 5 times. In the Assay Group, mice were given 2 mg/kg (in LABRAFIL M2125 CS, CAS:61789-25-1, Gattefosse, France) orally and rectally every day except weekends. Control Groups were given the same vehicle formulation. After the experiment reached 5 cycles of the DSS/non-DSS treatment, colons were collected and stained briefly with Alician Blue (1% in 3% Acetic Acid Solution) and rinsed in PBS for visualizing tumors. Colon images were analyzed by ImageJ for quantifying tumor size and number. Colon samples were subjected to paraffin embed for HE or Multiplex Immunofluorescence Staining. The mice were not randomized but were randomly allotted to the treatment groups. No a priori sample size estimated were performed, and the samples were chosen based on prior published sample sizes for treatment studies [43].

#### Metabolomics study

6-8 weeks FVB humanized PXR mice were divided into four groups, with 8 mice (n=8; 5 males and 3 females) in each experimental group. Mice had free access to water, containing 3% (w/v) DSS, and they were administered *quaque die* / *per os* + *per rectum* for 5 days with vehicle or 0.4 mg/kg FKK6. The feces were collected and subjected to metabolomic analyses.

### Multiplex Immunofluorescence staining

Immunofluorescence staining was performed by iHisto Inc. (ihisto.io) Paraffin sections were then deparaffinized and hydrated using the following steps: 15 min in xylene twice; 5, 5, and 5 min in 100%, 100%, and 75% ethanol, respectively; and 5 min in PBS at room temperature repeated three times. Antigen retrieval was achieved by heating the slides in a Citrate-based solution (VectorLab, H-3300) for 15 min in a pressure cooker (SB Bio) oven and 20 min of cooling at room temperature. Sections were washed with PBS three times and treated with 3% H2O2 for 15 min and 3% bovine serum albumin for 30 min. The sections were incubated with the primary antibody Rabbit Anti-Ki67 (abcam ab16667, 1:200) overnight at 4°C. Sections were rinsed with PBS and incubated with secondary antibody HRP-conjugated Goat anti-rabbit secondary antibodies (Invitrogen, 31460, 1:500) for 1 hour at room temperature. After rinsing with PBS, sections were incubated for 10 minutes at room temperature in Alexa Fluor™ 555 Tyramide Reagent (Invitrogen, B40955). Antigen retrieval was achieved by heating the slides in Citrate-based solution (VectorLab, H-3300) for 15 min in a pressure cooker (SB Bio) oven and 20 min of cooling at room temperature. Sections were washed with PBS three times, treated with 3% bovine serum albumin for 30 min. The sections were incubated with the primary antibody c-Myc (Cell Signaling 13987, 1:1000) overnight at 4°C. Sections were rinsed with PBS and incubated with secondary antibody HRP-conjugated Goat anti-rabbit secondary antibodies (Invitrogen, 31460, 1:500) for 1 hour at room temperature. After rinsing with PBS, sections were incubated for 10 minutes at room temperature in Alexa Fluor™ 488 Tyramide Reagent (Invitrogen, B40953). Antigen retrieval was achieved by heating the slides in Citrate-based solution (VectorLab, H-3300) for 15 min in a pressure cooker (SB Bio) oven and 20 min of cooling at room temperature. Sections were washed with PBS three times, treated with 3% bovine serum albumin for 30 min. The sections were incubated with Rabbit anti-cyclin D1 (Cell signaling 2978, 1:500) overnight at 4 °C. Sections were rinsed with PBS and incubated with secondary antibody HRP-conjugated Goat anti-rabbit secondary antibodies (Invitrogen, 31460, 1:500) for 1 hour at room temperature. After rinsing with PBS, sections were incubated for 10 minutes at room temperature in Alexa Fluor™ 594 Tyramide Reagent (Invitrogen, B40957). Antigen retrieval was achieved by heating the slides in a Citrate-based solution (VectorLab, H-3300) for 15 min in a pressure cooker (SB Bio) oven and 20 min of cooling at room temperature. Sections were washed with PBS three times and treated with 3% bovine serum albumin for 30 min. The sections were incubated with Rabbit anti-β-Catenin (Cell signaling 8480, 1:200) & Phospho-Histone H2A.X (eBioscience 14-9865-82, 1:100) overnight at 4 °C. Subsequently, the sections were immunohistochemically stained with Goat anti-Rabbit AF647 (Invitrogen A-21245, 1:500) & Goat anti-Mouse AF750 (Invitrogen A-21037, 1:500) for 1 hour at room temperature. Quench autofluorescence with 0.1% Sudan Black B for 15 min and stain DAPI. Whole slide scanning (40x) was performed on a Panoramic midi scanner (3D histech).

*Analysis of Multiplex Immunofluorescence staining:* The selected markers, paired with their corresponding fluorophores, comprised FITC for c-MYC, CY3 for Ki-67, CY5 for β-catenin, and Texas Red for cyclin D. Tissue sections labeled with immunofluorescence were visualized and captured using a 3D Histech P250 high-capacity slide scanner. The co-expressed staining within the delineated regions of interest (ROIs) was quantified using the HistoQuant Quant Center module (3DHISTECH) and was subsequently processed utilizing 3DHistech SlideViewer software, version 2.7. Out of n = 6 in each group, three randomly selected slides from each experimental group were excluded from the data analysis due to predefined exclusion criteria, such as suboptimal tissue preparation or unsuccessful immunostaining.

## RESULTS

### *In vitro* safety of FKK6

In addition to recently reported *in vitro* safety signals of FKK6 [36], we have performed mutagenic and genotoxic tests. An Ames fluctuation test was employed to assess the mutagenic potential of FKK6 across various strains of Salmonella typhimurium (TA98, TA100, TA1535, TA1537). The evaluation was carried out in the presence and absence of rat liver S9 fraction to detect potential metabolic activation. FKK6 did not induce an increase in revertant colonies in any tested strain (Figure 1A). FKK6 did not exhibit cytotoxic effects on the tested bacterial strains, implying that the lack of mutagenicity observed in the Ames test was not due to the growth inhibition of bacteria (Figure 1B). The genotoxic potential of FKK6 was assessed by the *in vitro* micronucleus test in CHO-K1 cells. FKK6 did not increase micronuclei formation, which supports its non-genotoxic profile (Figure 1C).

**Figure 1.**
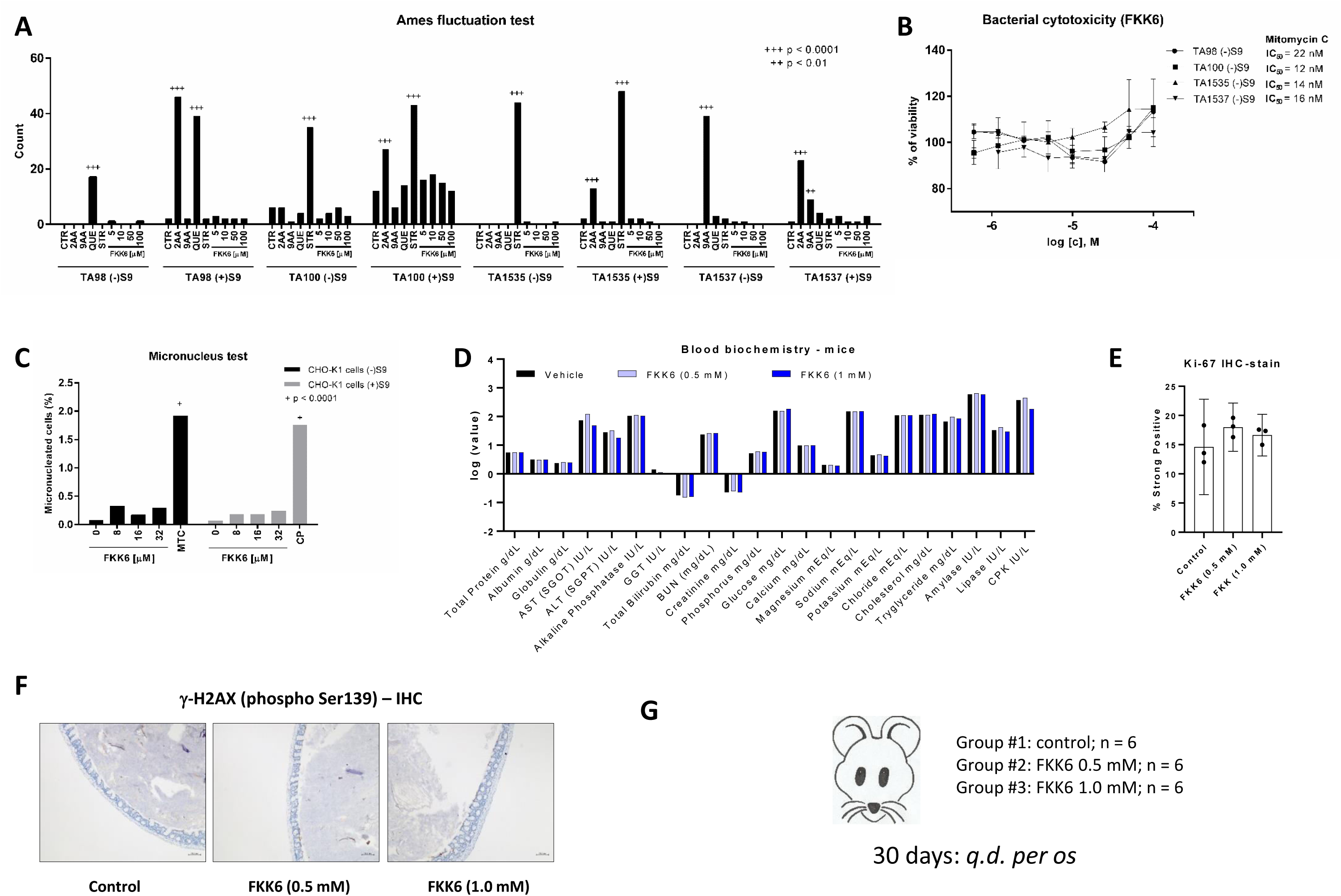
*In vitro* and *in vivo* safety of FKK6. ***(A,B) Ames fluctuation test & Bacterial cytotoxicity.*** Assays was carried out in four different strains of Salmonella typhimurium, in the presence or absence of rat liver S9 fraction, as described in detail in the Materials and Methods section. CTR = background, 2AA = 2-aminoanthracene (10 μM), 9AA = 9-aminoacridine (10 μM), QUE = quercetin (30 μM), STR = streptozotocin (2.5 μM); Significance was calculated using the one-tailed Fisher’s exact test. ***(C) Micronucleus assay.*** CHO-K1 cells were incubated at 37 °C with FKK6 (0 μM; 8 μM; 16 μM; 32 μM), Mitomycin C (MTC; 0.3 μM) or cyclophosphamide (CP; 72 μM) in the presence (4 h) or the absence (24 h) of rat liver S9 fraction. The percent of micronucleated cells was calculated and the significance was determined using the t-test. ***(D) Mouse blood biochemistry.*** 30-days chronic toxicity *in vivo* experiment; FKK6 (0 mM; 0.5 mM; 1 mM) was dosed *q.d./p.o.*; n = 6 mice *per* group. ***(E) Ki-67 intestinal expression - immunohistochemistry.*** Evaluated in n = 3 mice *per* group. ***(F) γ-H2AX (phospho Ser139) intestinal expression - immunohistochemistry.*** Evaluated in n = 3 mice *per* group. ***(G) Mice 30-days chronic toxicity experiment scheme***

### *In vivo* safety of FKK6

We evaluated the chronic toxicity of FKK6 in C57BL/6 mice (n=6; each group) administered FKK6 (1 mg/kg and 2 mg/kg in normal saline with 1% DMSO) via oral gavage daily for 30 days (Figure 1G).

Histological examination of mouse tissues revealed no pathological changes indicative of FKK6 toxicity (Table 1). Biochemical blood analyses showed no decline in FKK6-treated mice compared to vehicle-treated mice (Figure 1D). Immunohistochemical staining of intestines for Ki-67, a cell proliferation marker, did not show a difference between FKK6-treated and control mice, implying that FKK6 does not promote hyperplasia or neoplasia (Figure 1E). Similarly, staining of intestines for γ-H2AX, a marker of DNA damage, revealed FKK6 safety regarding genotoxicity (Figure 1F), which is consistent with in vitro data (Figure 1A-C).

**Table 1.**
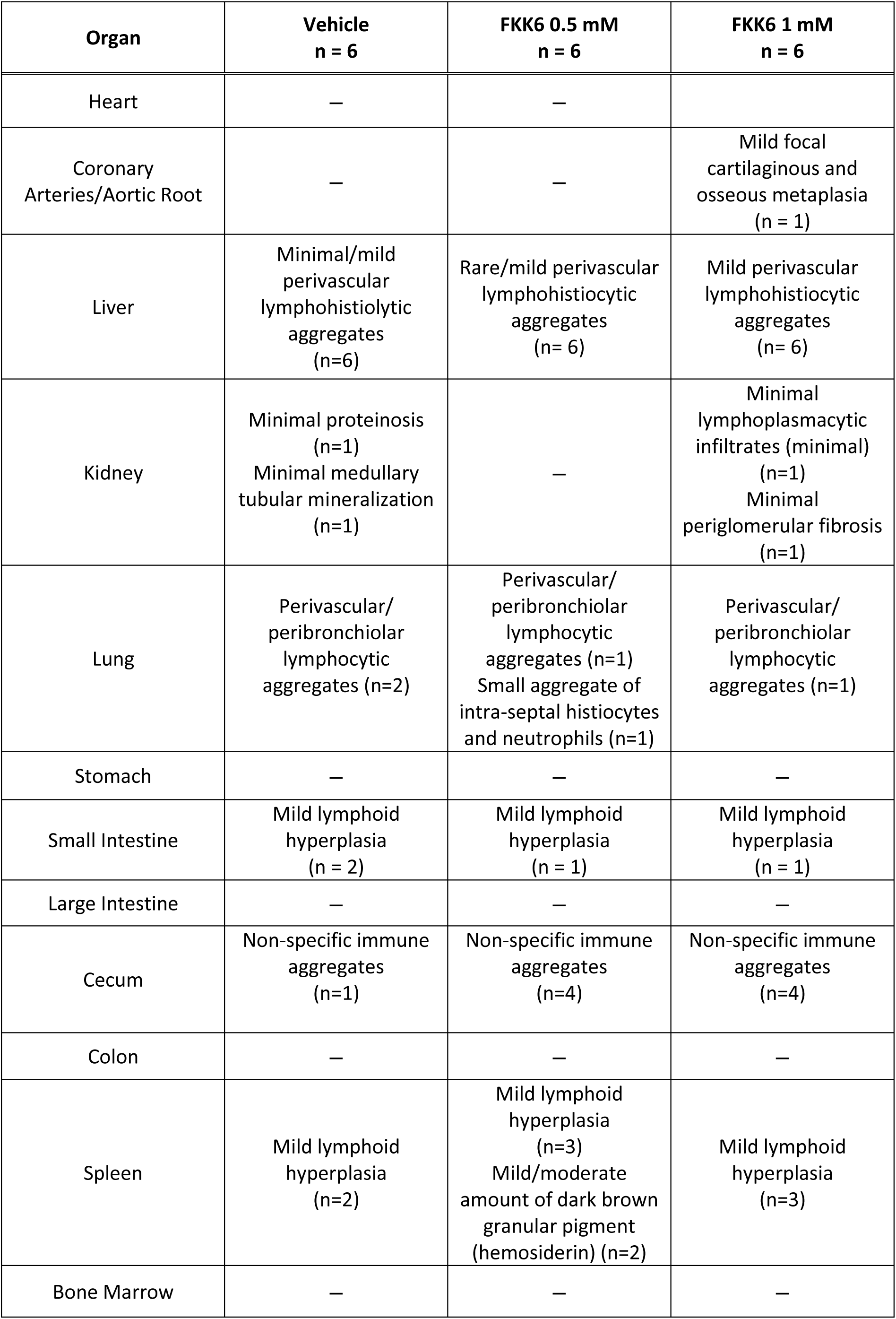
30-day chronic toxicity *in vivo* experiment – histology. : FKK6 (0 mM; 0.5 mM; 1 mM) was dosed *q.d./p.o.*; n = 6 mice *per* group.

### Comparative metabolomics of FKK6 in DSS-treated mice

In a previous study, we reported PXR-dependent protective effects of FKK6 in mice with dextran sodium sulfate (DSS) induced colitis [35]. Therefore, we conducted comparative fecal metabolomic analyses in mice exposed for 5 days to DSS and administered in parallel with FKK6 (0.4 mg/kg; qd/po+pr) or vehicle. The metabolic clusters, including intermediary metabolism of amino acids, fatty acids, nucleotides, carbohydrates, neurotransmitters, and vitamins, were not altered in FKK6-treated mice compared to the control group (Figure 2).

**Figure 2.**
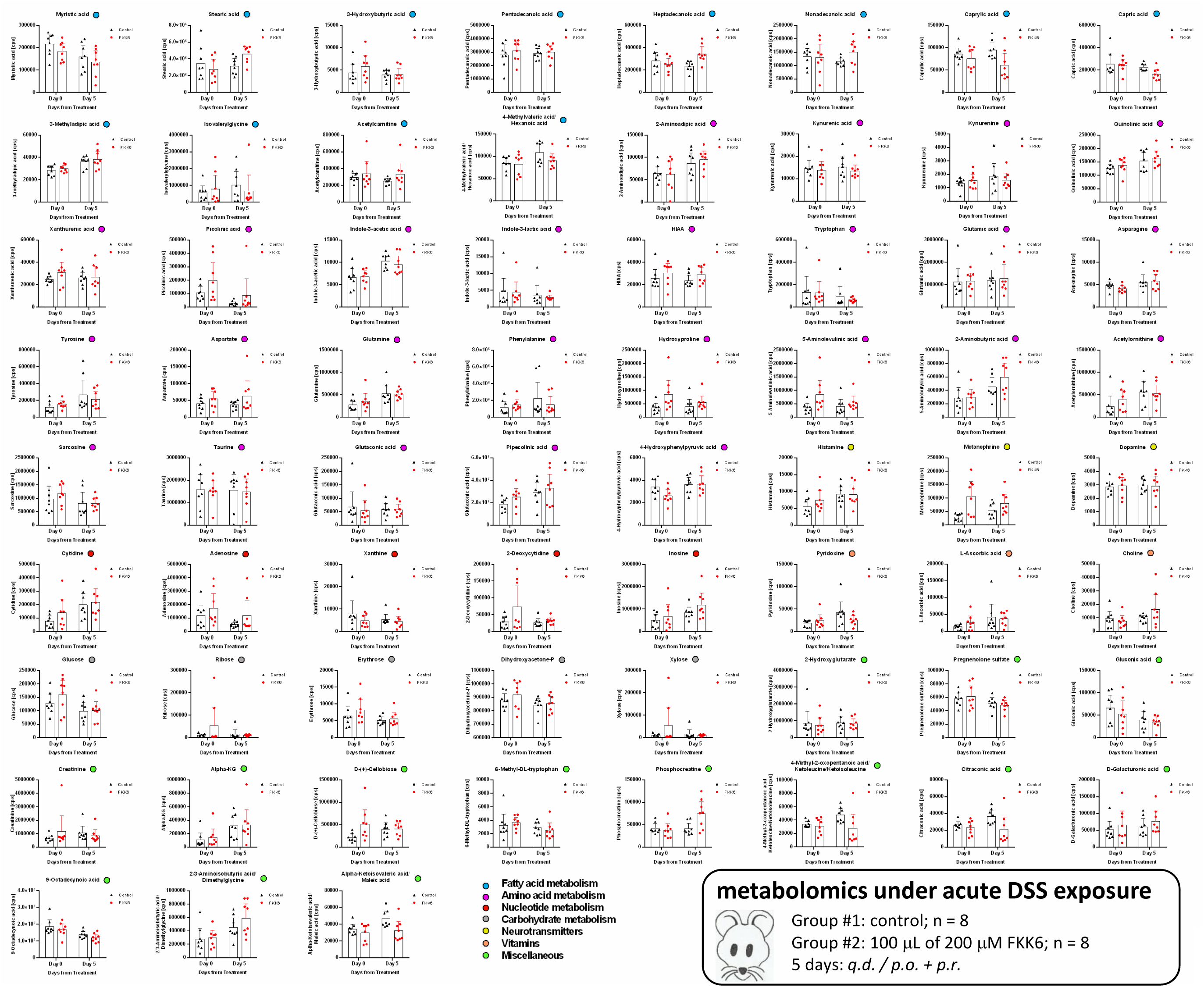
FKK6 does not alter the metabolomic profile of mouse feces. Comparative metabolomic analyses are shown. Mice were divided into four groups, with 8 mice (n=8; 5 males and 3 females) in each experimental group. Mice had free access to water, containing 3% DSS and were administered *quaque die* / *per os* + *per rectum* for 5 days with vehicle or 0.4 mg/kg FKK6. The data are expressed as mean ± SD.

Altogether, *in vitro* data, 30-day in vivo chronic toxicity assessment, and in vivo, comparative metabolomics, together with recently published pharmacological profiling [36], support FKK6 therapeutic drug safety.

### FKK6 exhibits anti-cancer activity in a mice model of AOM/DSS-induced colitis-associated colon cancer

Since FKK6 displayed anti-inflammatory activity *in vitro* and in mice with DSS-induced colitis [35], we investigated whether FKK6 is protective against AOM/DSS-induced colitis-associated intestinal tumorigenesis in mice. The experimental design of an AOM/DSS animal study is illustrated in Figure 3. Mice subjected to AOM/DSS protocol were divided into two groups; a control group receiving vehicle (n=8) and an FKK6-treated group (n=10) receiving every day (except weekends) 2 mg/kg FKK6 (po+pr). The number and size of tumors were evaluated after staining mouse colons with Alcian blue. Representative macroscopic views of the colon tumor area provide a direct visual comparison between the control and FKK6-treated groups. The sections of the colon from the control group display numerous and large tumors, as indicated by yellow circles (Figure 4B, Figure S1A). In contrast, in the colon sections from the FKK6-treated group, a reduction in tumor size and number is evident (Figure 4C, Figure S1B). Quantitative analysis using ImageJ software was consistent with visual observations. The scatter plots show a substantial and significant (p<0.01) decrease in the number and size of tumors in the FKK6-treated group (Figure 4A). Mouse colon tissues were subjected to immunohistochemical analyses. Colon sections were stained with hematoxylin and eosin (H&E) to evaluate overall tissue morphology and the extent of tumor invasion. Whereas the control group exhibited substantial disruption of architecture and invasive tumor growth, characteristic of advanced neoplastic transformation (Figure 5A, Figure S2A), the FKK6-treated group maintained more intact histological structure with minimal signs of invasion and disruption (Figure 5D, Figure S2B). In parallel, colonic sections were stained by multiplex immunofluorescence to assess the expression of key tumor markers, including c-MYC, β-catenin, Ki-67, and cyclin D. Fluorescence intensity quantification in the regions corresponding to tumors (Figure 5G) revealed an 85.1% reduction in c-MYC and an 82% reduction in combined markers (β-catenin, Ki-67, cyclin D) in the FKK6-treated group (Figure 5B-C) as compared to control group (Figure 5E-F). Collectively, FKK6 displayed substantial anti-tumor activity in the AOM/DSS mouse model of inflammation-related colorectal cancer, which is documented by reduced size and number of tumors and decreased expression of tumor markers.

**Figure 3.**
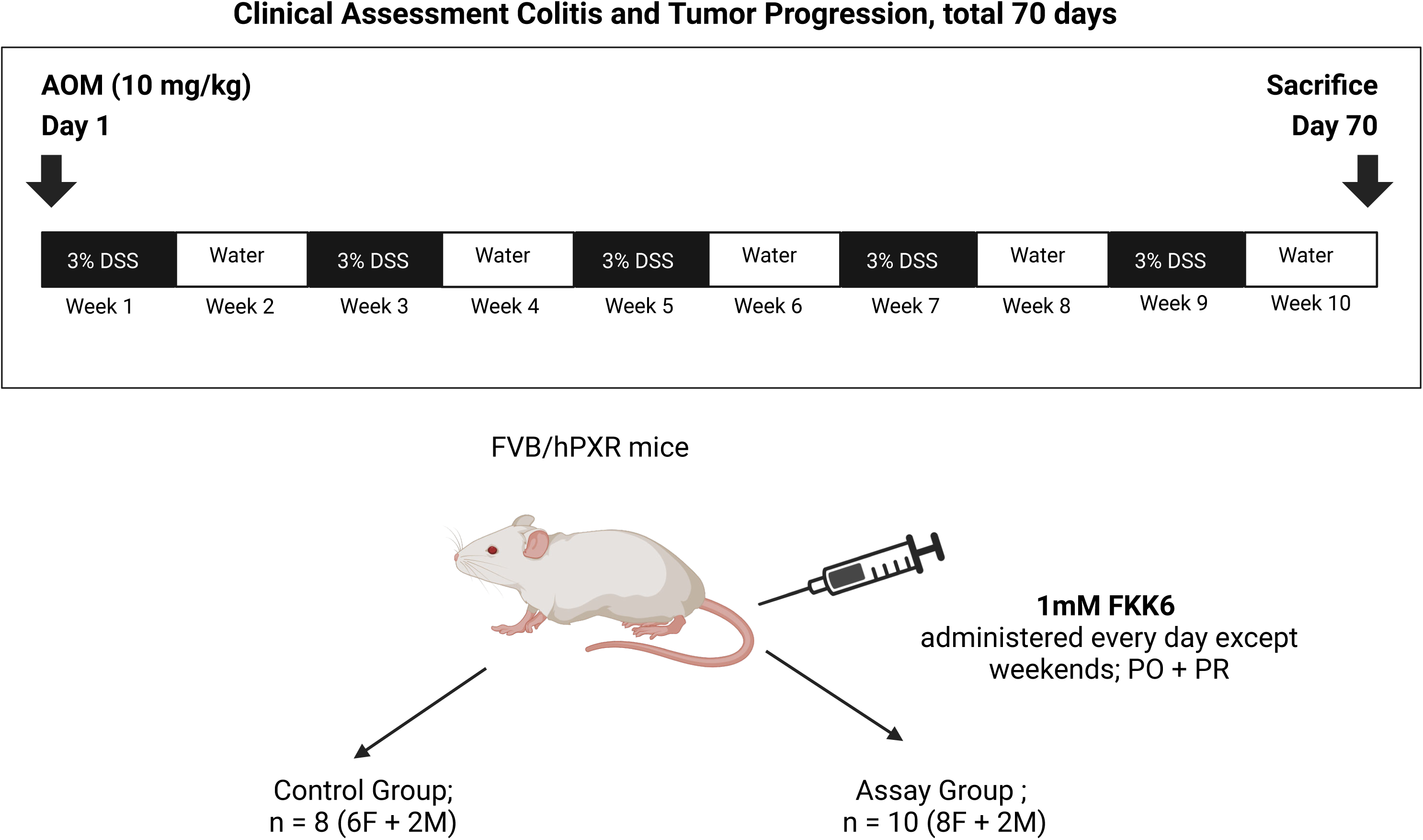
Schematic overview of AOM/DSS animal study design.

**Figure 4:**
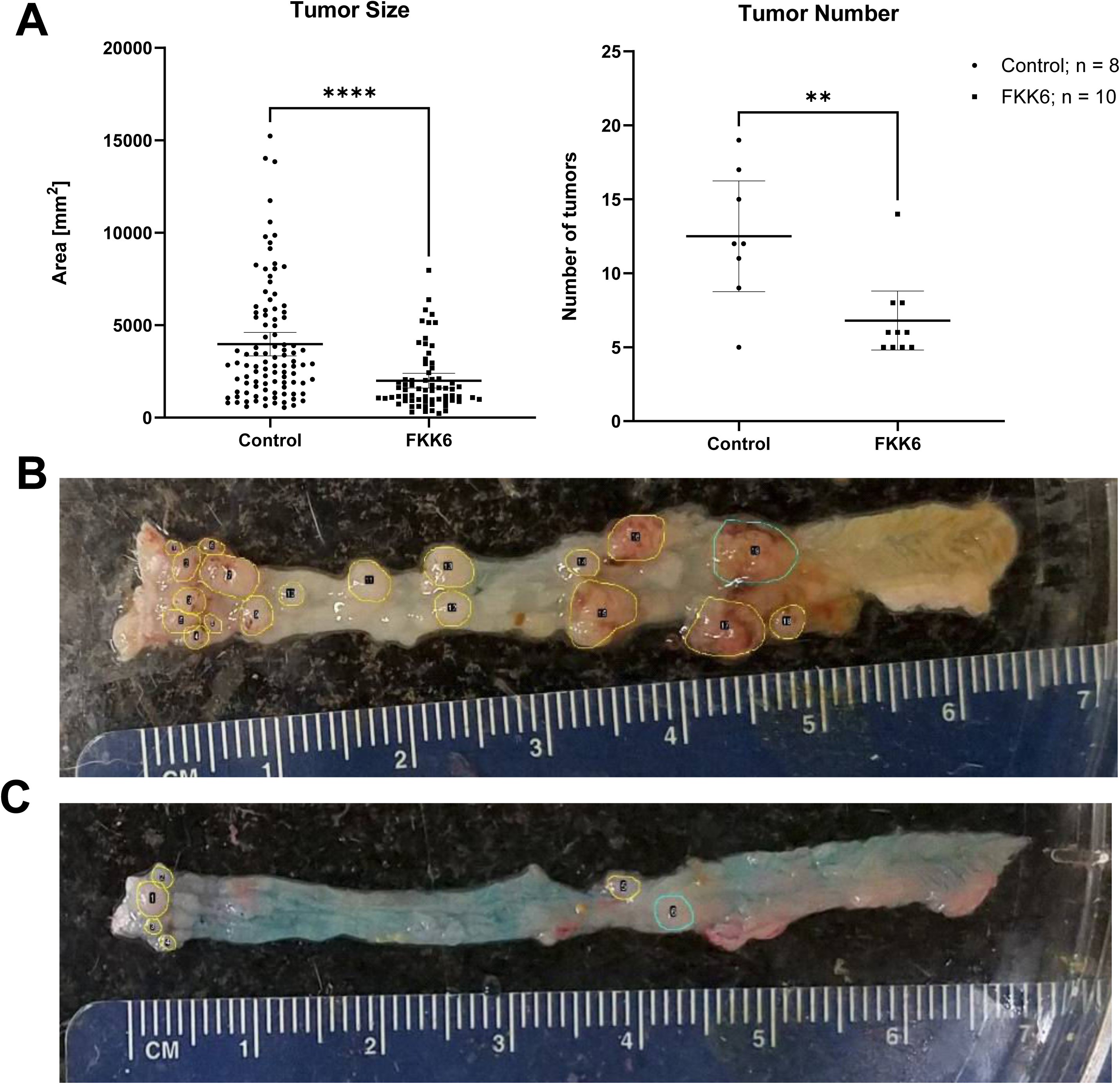
FKK6 exhibits protective effects in the AOM/DSS mouse model *in vivo*. ***(A)*** *The size of tumors* (left plot): each dot (circle, square) represents an individual tumor, and a plot comprises cumulatively all tumors in the examined colonic section of all mice in the experimental group. *The number of tumors* (right plot): each dot (circle, square) represents a number of tumors in the examined colonic section of individual mice in the experimental group. Statistical analysis was conducted using the Mann-Whitney test, **, P < 0.01; ***(B)*** A representative macroscopic view of the colonic section of control mouse (n = 8); ***(C)*** A representative macroscopic view of the colonic section of FKK6-treated mouse (n = 10). Tumor quantification was performed using the ImageJ software.

**Figure 5:**
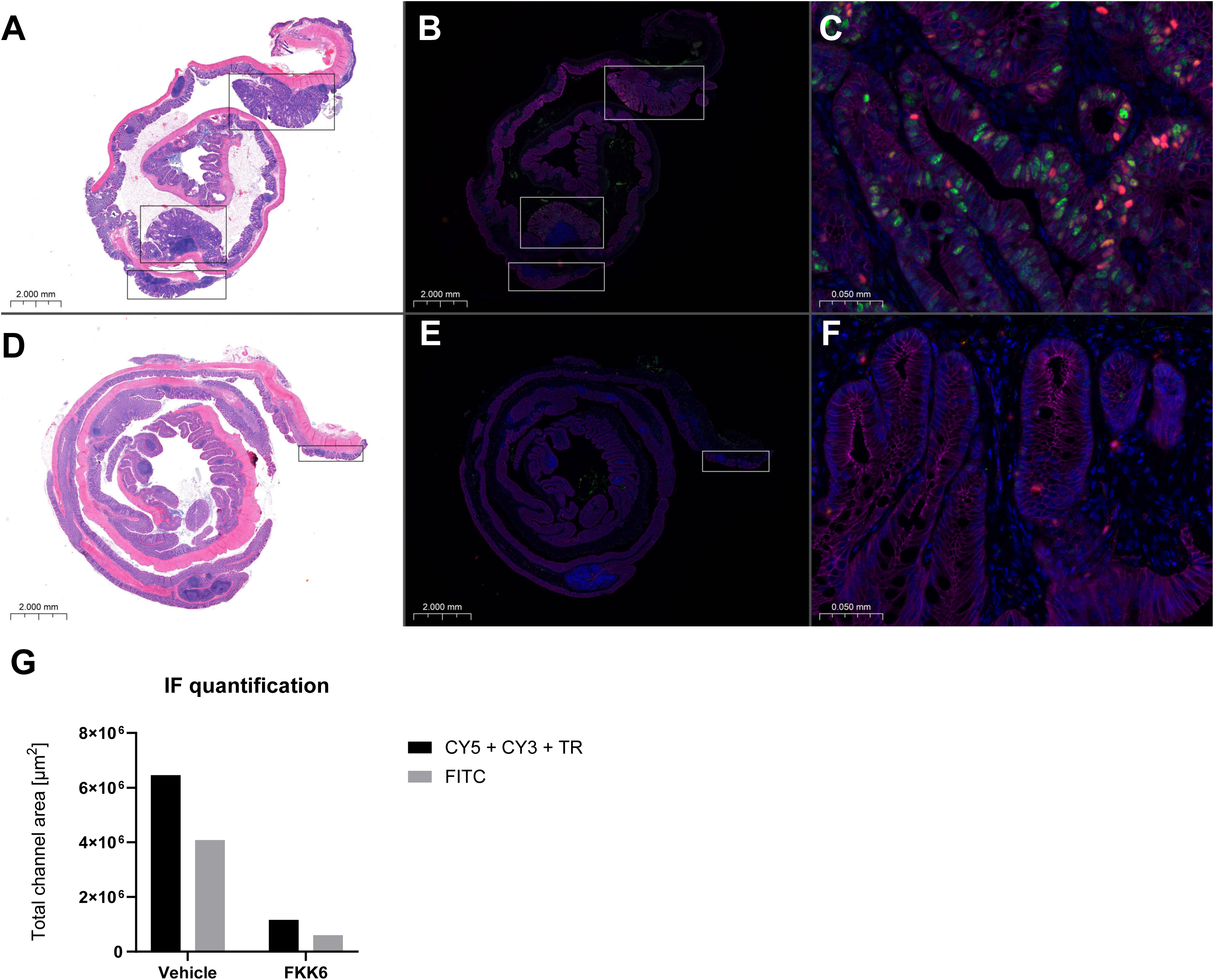
FKK6 reduces tumor markers in the colon of AOM/DSS-treated mice. Representative images from the control group ***(A, B, C)*** and the FKK6-treated group ***(D, E, F)*** are shown. Mouse colon sections were counterstained with H&E (***A,D***), and the tumor area was indicated by an inserted black frame. The same colon sections were immune-fluorescently labeled with anti-c-MYC (FITC), anti-Ki-67 (CY3), anti-β-catenin (CY5), and anti-cyclin D (Texas Red) antibodies (***B,E*** – low magnification; ***C,F*** – high magnification). The tumor markers in the areas indicated by the white frame (***B, E***) were quantified (n = 3; for each group), and a detailed view of these areas is presented (***C, F***). Immunohistochemical quantification of tumor markers was performed using 3DHistech SlideViewer software. The results are presented as absolute values corresponding to individual channels in tumor area (***G***).

## DISCUSSION

The development of FKK6 as a potential therapeutic agent for inflammatory bowel disease (IBD) and colorectal cancer (CRC) represents a significant advancement in harnessing microbial metabolite mimicry for drug discovery. The AOM/DSS model faithfully mimics the pathogenesis of human CAC, characterized by chronic inflammation leading to dysplasia and carcinoma [44]. The use of FKK6 in this model is particularly compelling. The anti-inflammatory and anti-tumor effects of FKK6 could translate into significant therapeutic benefits for patients with IBD and those at risk of developing CAC. By targeting PXR, FKK6 addresses the inflammatory component of these diseases and offers a novel approach to cancer prevention and treatment.

The comprehensive genotoxicity and toxicity testing results suggest that FKK6 does not exhibit mutagenic potential or significant cytotoxic effects. Evaluation of compound chronic toxicity over 30-days in mice also supported its safety profile, with no significant histopathological changes observed in major organs. While mild immune reactions were noted in the liver, lungs, and spleen, further investigation is required to understand the immunomodulatory effects of FKK6 fully.

Safety assessment of FKK6 *in vitro*, as detailed in the safety profile study [36], revealed several key pharmacological properties supporting its potential for clinical development. FKK6 exhibited high permeability, solubility, and metabolic clearance, suggesting favorable pharmacokinetics. The compound was metabolized primarily through reduction and oxidation reactions, but high clearance may limit its oral bioavailability. Interestingly, we assessed the direct effects of FKK6 on fecal metabolomics on day 5 of treatment with FKK6 in 3% DSS exposed mice. Notably, mice have preserved weight by day 5 and only after day 5 does the weight begin to decline, so at day 5 host effects on fecal metabolomes are thought to be much less than if we assessed at a point when the weights decline. In this assessment, FKK6 had minimal effects on various metabolites belonging to different classes. While every metabolite was not assessed broadly, FKK6 has minimal effects on different metabolites, suggesting minimal off-target effects in this context.

Based on the safety profile and therapeutic potential of FKK6, several promising clinical applications can be envisioned. However, optimization of FKK6’s biological availability through the development of enhanced formulations is crucial for its clinical translation. Oral delivery system development is currently an active area of research in treating diseases affecting the colon. Nanotechnology has recently emerged as a new and effective tool for targeted drug delivery. For example, as demonstrated by Merlin’s group [45, 46], encapsulation of drugs in lipid nanoparticles significantly improves their pharmacokinetic properties compared to free compounds. This means oral nanoparticle-based delivery systems may protect the charged drug from the harsh gastrointestinal environment and selectively increase drug concentration at the disease site. Therefore, considering the compound FKK6, investigating oral formulations that ensure safe and effective delivery to the colon may be beneficial in maximizing anti-inflammatory effects while minimizing systemic exposure.

In the context of anti-tumor effects, treatment with the tested compound significantly reduced the number and size of tumors compared to the control group. Histological and immunohistochemical analyses showed that FKK6 treatment maintained a more normalized tissue structure and reduced the expression of key tumor markers. These results indicate that FKK6 acts as an anti-tumor by modulating critical oncogenic pathways. Given the complexity of cancer pathogenesis, the mechanism of FKK6 action could also involve influencing the tumor microenvironment and inflammatory drivers of colon cancer.

The comprehensive evaluation of FKK6 has demonstrated its favorable safety profile and strong anti-tumor effects in colorectal cancer, potentially by modulating critical oncogenic pathways. The absence of mutagenic and cytotoxic effects and the lack of significant adverse effects in chronic toxicity studies underscores the compound’s safety. The significant reduction in tumor burden and modulation of oncogenic pathways further highlight the therapeutic potential of FKK6 in colorectal cancer.

## STUDY LIMITATIONS

The dosing of FKK6 in the DMSO formulation has not been optimized for mice so lower or even higher doses tolerated by mice could potentially have a greater effect on reducing tumor burden. Other formulations of FKK6, currently under development, have not been tested in the mouse model. The full pharmacokinetic profile of single and multi-dosing of FKK6 has not be characterized and in the future these data may help guide optimal dosing and delivery. A mechanistic evaluation of FKK6 would be important to know how FKK6 might prevent tumor formation and there would need to be cautious application of FKK6 when tumors are already present [15]. Additional models of inflammation-induced colon cancer also need to be studied in the context of FKK6 effects. The fecal metabolomics also need to be expanded to other unstudied metabolites and on different intestinal inflammation and cancer models.

## ACKNOWLEDGEMENTS

Financial support from NIH #R01CA222469 (to S.M) and SIG #1S10OD026852-01A1 (to H.G.), Czech Science Foundation grant #22-00355S (to Z.D.), Albert Einstein Cancer Center Support Grant of the National Institutes of Health #P30CA013330 (to A.P.B.), and National Cancer Institute grant #P30CA013330 (to H.Guzik) is greatly acknowledged.

## AUTHORŚ CONTRIBUTIONS

CrediT STATEMENT: Conceptualization**^1^**, Data Curation**^2^**, Formal Analysis**^3^**, Funding Acquisition**^4^**, Investigation**^5^**, Methodology**^6^**, Project Administration**^7^**, Resources**^8^**, Supervision**^9^**, Validation**^10^**, Visualization**^11^**, Writing – Original Draft Preparation**^12^**, Writing – Review & Editing**^13^** Lucia Sládeková^2,3,5,6^, Hao Li^2,3,5,6^, Vera M. DesMarais^2,3,5,6^, Amanda P. Beck^2,3,5,6^, Hillary Guzik^2,3,5,6^, Barbora Vyhlídalová^11^, Haiwei Gu^2,3,5,6,8^, Sridhar Mani^1,2,3,4,5,7,8,9,10,12,13^, Zdeněk Dvořák^1,2,3,4,5,7,8,9,10,12,13^

## DATA AVAILABILITY STATEMENT

The data supporting this study’s findings are openly available in ZENODO at http://doi.org/10.5281/zenodo.12187327.

## ABBREVIATIONS

AOM: azoxymethane
CAC: colitis-associated colon cancer
COX-1/2: cyclooxygenase 1/2
CRC: colorectal cancer
DSS: dextran sodium sulfate
IBD: inflammatory bowel disease
NSAID: non-steroidal anti-inflammatory drug
PXR: pregnane X receptor
UC: ulcerative colitis

---

**Figure S1:**
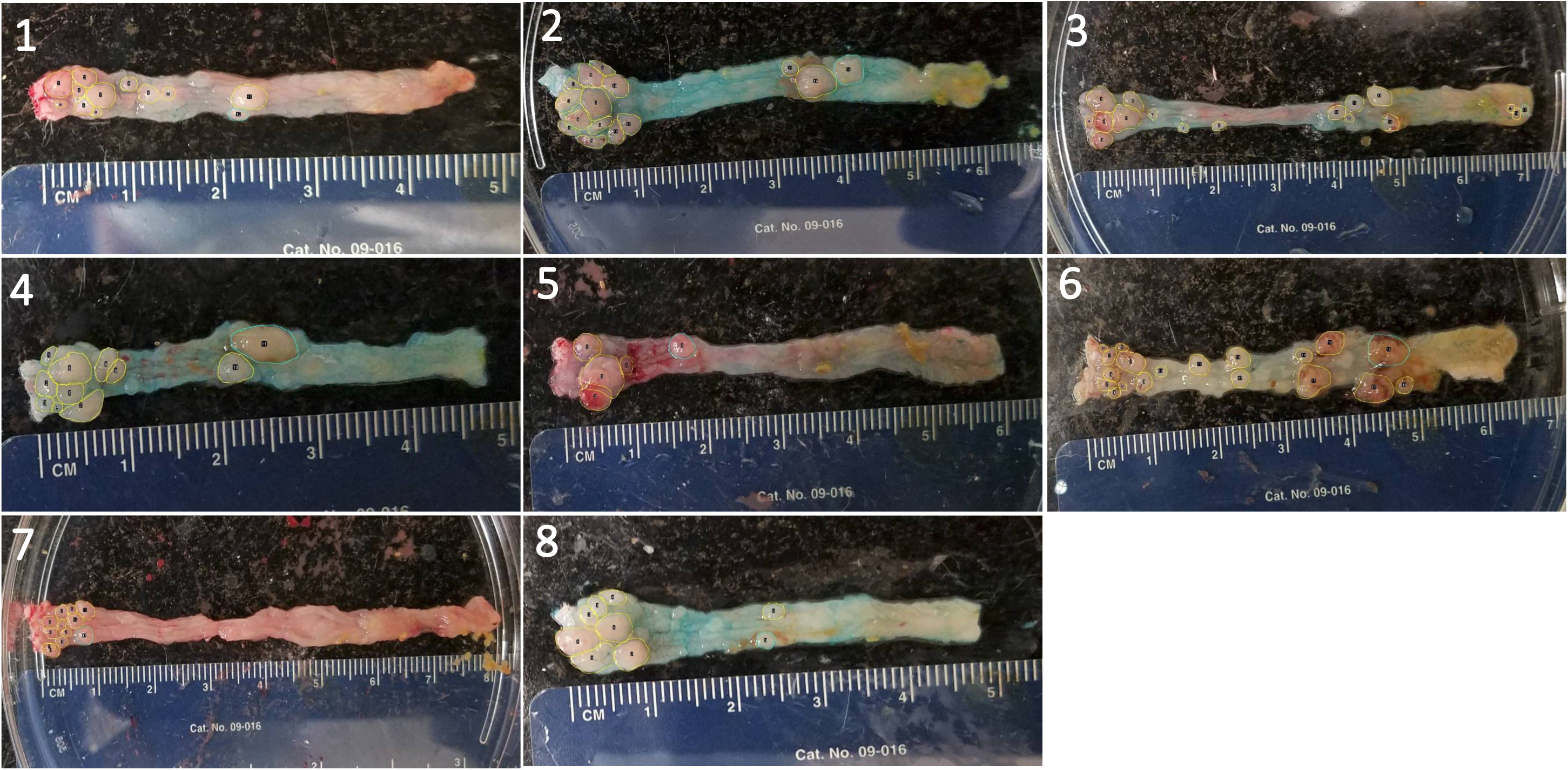
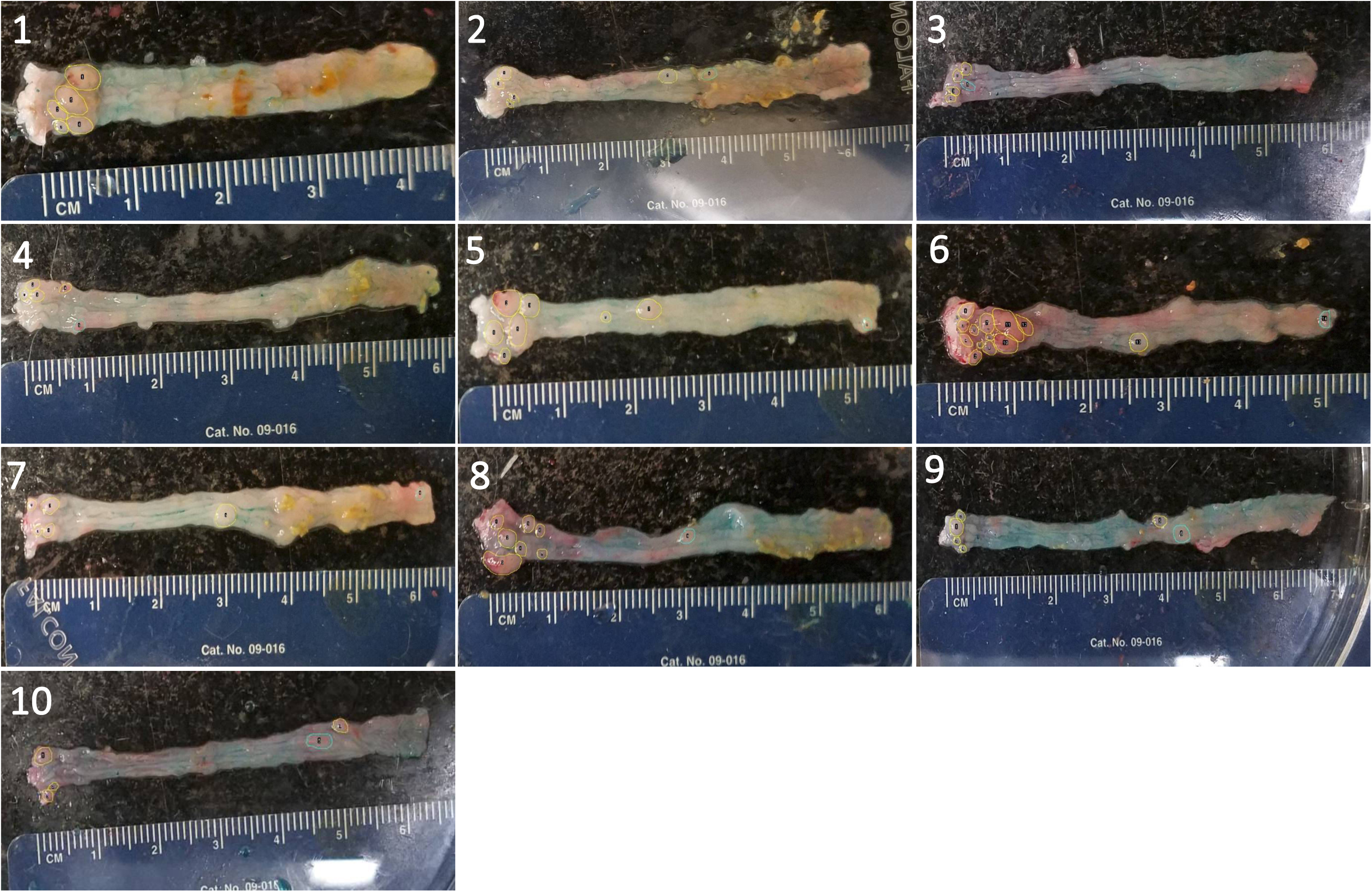
Mice colonic tumor visualization by Alician blue. ***(A)*** Control group (n = 8). ***(B)*** FKK6-treated group (n = 10).

**Figure S2:**
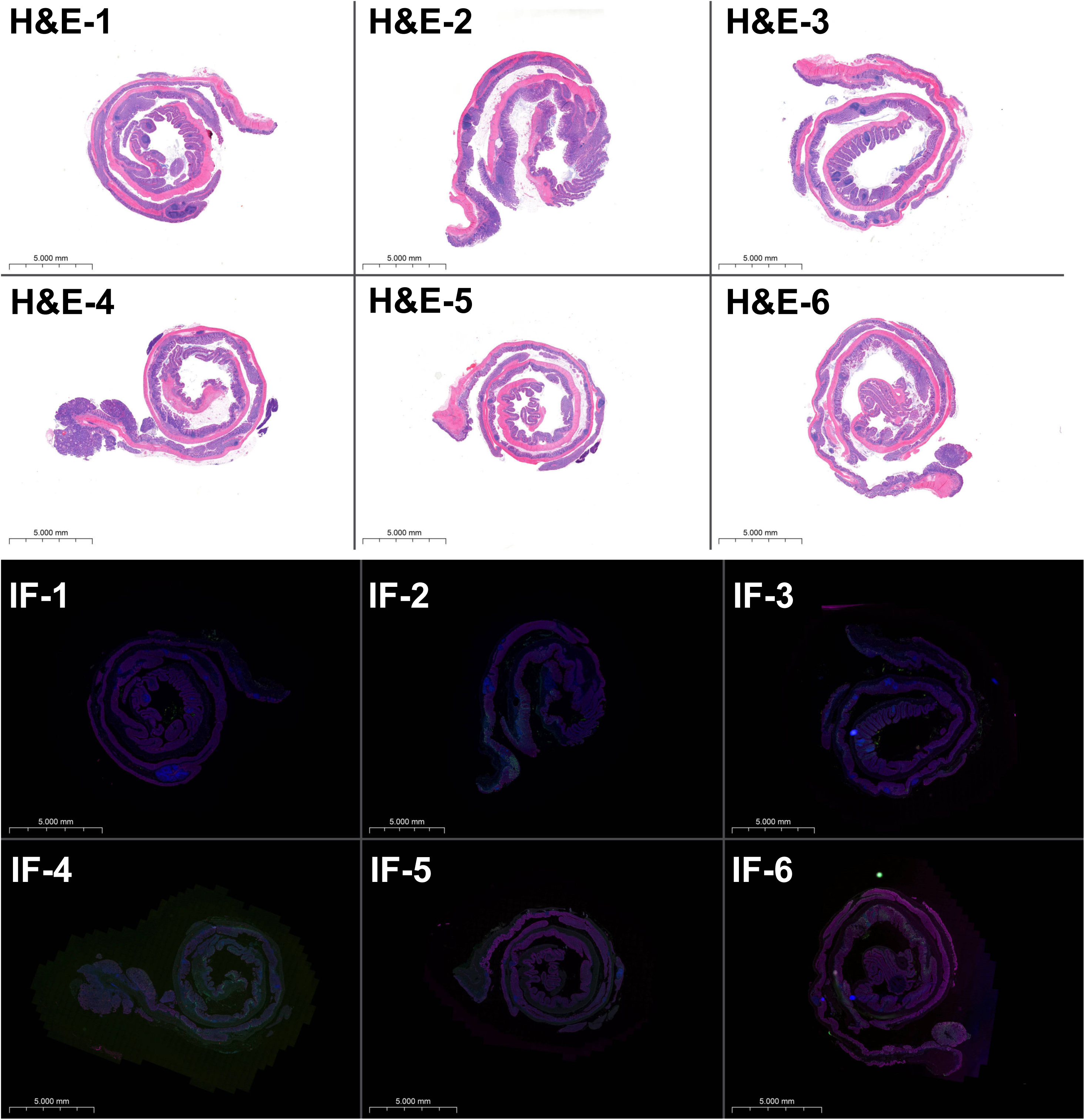
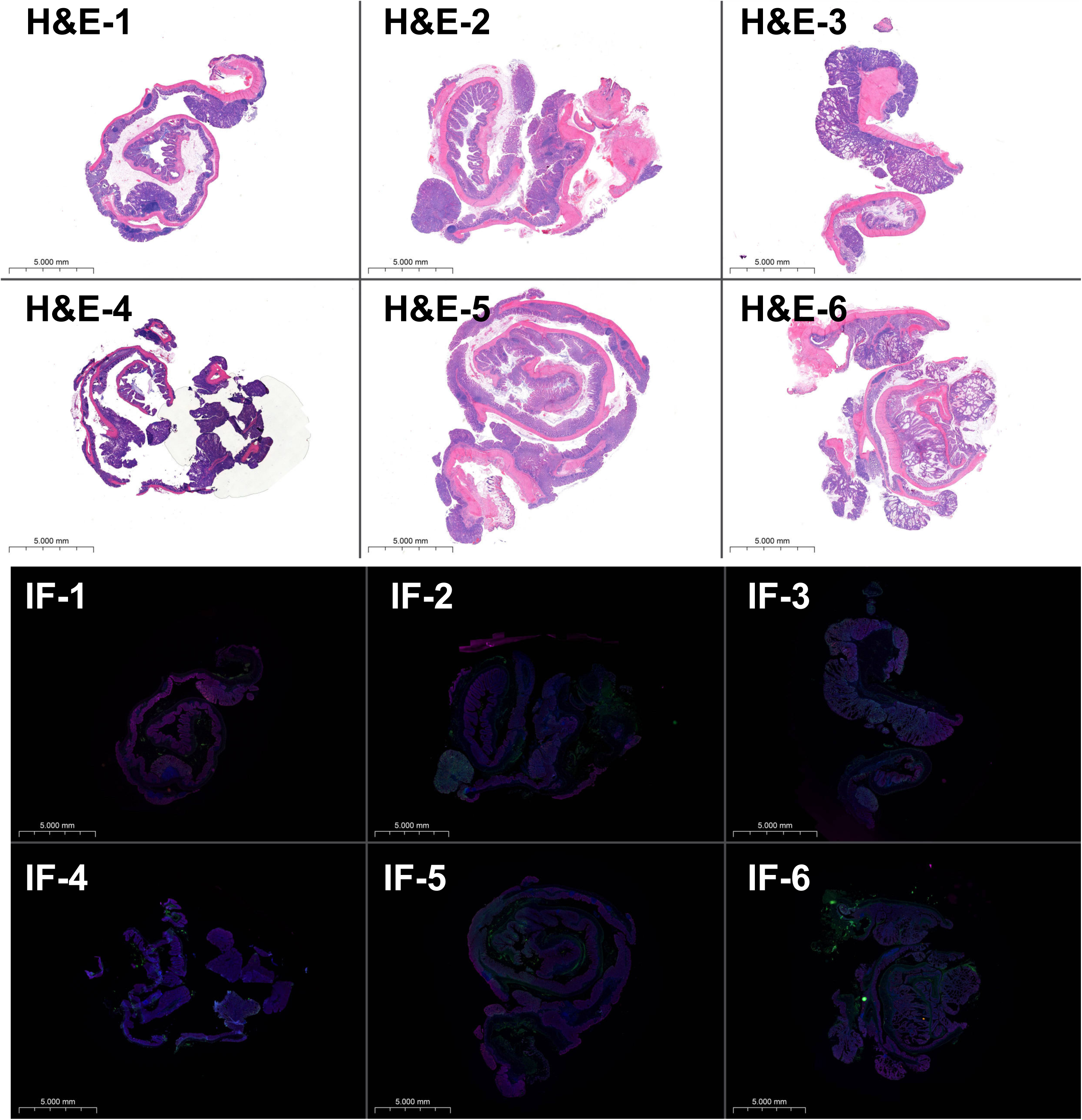
Mice colonic tissue H&E and IF staining. Mouse colon sections were stained with H&E, and immuno-fluorescently labeled with anti-c-MYC (FITC), anti-Ki-67 (CY3), anti-β-catenin (CY5), and anti-cyclin D (Texas Red) antibodies. ***(A)*** control (AOM/DSS) group (n=6). ***(B)*** FKK6 (+AOM/DSS) group (n=6). Note: Samples 4, 5, and 6 were excluded from IF quantification due to suboptimal preparation or staining.

